## Supplementary Figures and Tables for "T Cell-to-Stroma Enrichment (TSE) score: a gene expression metric that predicts response to immune checkpoint inhibitors in patients with urothelial cancer"

**Supplementary Table 1. Characteristics of 70 patients with metastatic urothelial cancer who received treatment with pembrolizumab, and stratified by response to treatment at 6 months according to RECIST v1.1.**

| Cohort | Responder | Non-responder | Association with | | |
|---|---|---|---|---|---|
| **n=70** | n=24 | n=46 | Response | OS | PFS |
| Age in years: median (range) | 72 (42-85) | 69 (48-83) | p=0.15 | p=0.09[a] | p=0.09[a] |
| Male sex in numbers (%) | 22 (92%) | 36 (78%) | p=0.20 | p=0.045 | p=0.16 |
| Primary tumor location | | | | | |
| Bladder | 19 (79%) | 35 (76%) | p=1.0 | p=0.61[b] | p=0.79[b] |
| Upper tract | 5 (21%) | 10 (22%) | p=1.0 | | |
| Unknown | 0 | 1 (2%) | | | |
| Systemic pretreatment | | | | | |
| Yes | 18 (75%) | 45 (98%) | p=0.005 | p=0.026 | p=0.005 |
| Gemcitabine, cisplatin and/or carboplatin | 16 (67%) | 39 (85%) | | | |
| Methotrexate, vinblastine, doxorubicin, cisplatin | 2 (8%) | 4 (9%) | | | |
| Chemotherapy in combination with immunotherapy | 0 | 1 (2%)[c] | | | |
| Other[d] | 0 | 1 (2%) | | | |
| Radiotherapy pretreatment | | | | | |
| Yes | 2 (8%) | 15 (33%) | p=0.04 | p=0.047 | p=0.045 |
| No | 22 (92%) | 31 (67%) | | | |
| Best overall response (RECIST v1.1) | | | | | |
| Complete response | 3 (13%) | 0 | | p<0.001 | p<0.001 |
| Partial response | 18 (75%) | 1 (2%) | | | |
| Stable disease | 3 (13%) | 7 (15%) | | | |
| Progressive disease | 0 | 38 (83%) | | | |
| PD-L1 Combined positivity score | | | | | |
| Positive | 5 (21%) | 11 (24%) | p=1.0 | p=0.31[b] | p=0.47[b] |
| Negative | 8 (33%) | 16 (35%) | | | |
| Unknown | 11 (46%) | 19 (41%) | | | |
| Biopsy sites | | | | | |
| Lymph node | 14 (58%) | 17 (37%) | p=0.13 | p=0.14[f] | p=0.092[f] |
| Liver | 3 (13%) | 11 (24%) | p=0.35 | | |
| Soft tissue | 3 (13%) | 7 (15%) | p=1.0 | | |
| Other[e] | 4 (17%) | 11 (24%) | | | |
| RNA sequencing data available | 13 (54%) | 28 (61%) | | | |
| Immunofluorescence staining | 7 (35%) | 13 (65%) | | | |

[a]Age was dichotomized into young (age < median) and old.
[b]Unknown was excluded.
[c]Patient was randomized for nivolumab or placebo in trial.
[d]Other systemic pretreatments include a randomized trial treatment with adjuvant nivolumab or placebo treatment.
[e]Other biopsy sites include lung, bone, primary tumor, local recurrence, adrenal gland, and peritoneum (see also Fig 1).
[f]Others were excluded.

**Supplementary Table 2. Gene expression signatures for T cells, other immune cells (non-T cells) and stromal resident cells and products.**

| **T cells and products (n=21)** | Individual signatures |
| --- | --- |
| Ayers et al.[1] | T cell inflammation gene expression profile (GEP), Immune gene signature, interferon (IFN) gamma |
| Hammerl et al.[2] | Co-inhibitory receptors, co-stimulatory receptors, chemo-attractants, immune effector cells, adhesion molecules |
| Jerby-Arnon et al.[3] | T helper type 1 cells (Th1), T helper type 2 cells (Th2), T helper type 9 cells (Th9), T helper type 17 cells (Th17), T helper type 22 cells (Th22), T follicular helper cell (Tfh), naïve CD4 T cells, regulatory T cells (Treg), cytotoxic CD8 T cells, naïve T cells |
| Mariathasan et al.[4] | Eight-gene cytotoxic T cell transcriptional signature (tGE8) |
| Oh et al.[5] | Proliferating cytotoxic CD4 T cells |
| Spranger et al.[6] | T cell signature |
| **Immune cells (non-T cells; n=7)** | |
| Jerby-Arnon et al.[3] | B cells, natural killer (NK) cells, myeloid derived suppressor cells, plasmacytoid dendritic cells, myeloid dendritic cells, neutrophil, macrophages |
| **Stromal resident cells and their products (n=7)** | |
| Hammerl et al.[2] | Fibroblasts, endothelial cells, transforming growth factor beta (TGFb) |
| Jerby-Arnon et al.[3] | Cancer associated fibroblasts (CAF) |
| Mariathasan et al.[4] | TGF-β-induced genes (pan-TGF-β responsive signature: TBRS) |
| Wang et al.[7] | EMT/stroma core genes |
| Yoshihara et al.[8] | Stromal signature |

The complete list of genes per gene expression signature is available in Supplementary Data 3.

**Supplementary Table 3. Area under the curve (AUC) of receiver operating characteristic curves for predicting response to pembrolizumab with gene expression signatures.**

| Category | Name | AUC | 95% CI | Adj. p-value |
|---|---|---|---|---|
| Combined signatures | | | | |
| | TSE score | 0.88 | 0.78-0.99 | |
| | T cells | 0.75 | 0.58-0.92 | |
| | Stromal cells | 0.73 | 0.57-0.90 | |
| | Other immune cells (non T cells) | 0.62 | 0.40-0.84 | |
| | T cells + Other immune cells | 0.69 | 0.50-0.89 | |
| Individual signatures | | | | |
| T cells | IFN gamma | 0.77 | 0.61-0.93 | 0.052 |
| Stromal cells | TBRS | 0.77 | 0.61-0.92 | 0.038 |
| T cells | Cytotoxic CD8 T cells | 0.76 | 0.60-0.92 | 0.049 |
| T cells | tGE8 | 0.76 | 0.59-0.93 | 0.049 |
| T cells | Th9 | 0.76 | 0.59-0.93 | 0.083 |
| T cells | T cell inflamed GEP | 0.76 | 0.59-0.92 | 0.052 |
| Stromal cells | Fibroblasts | 0.76 | 0.60-0.91 | 0.049 |
| T cells | Th22 | 0.74 | 0.58-0.91 | 0.049 |
| T cells | Naive CD4 T cells | 0.74 | 0.55-0.93 | 0.067 |
| T cells | Immune gene signature | 0.73 | 0.55-0.91 | 0.040 |
| T cells | Th2 | 0.73 | 0.54-0.91 | 0.074 |
| Stromal cells | TGFb | 0.73 | 0.57-0.88 | 0.035 |
| Stromal cells | EMT/stroma core genes | 0.72 | 0.56-0.88 | 0.038 |
| T cells | T cell signature | 0.72 | 0.53-0.90 | 0.040 |
| T cells | Tfh | 0.72 | 0.52-0.91 | 0.047 |
| T cells | Chemoattractants | 0.71 | 0.52-0.91 | 0.040 |
| Stromal cells | CAF | 0.71 | 0.53-0.88 | 0.035 |
| T cells | Co-stimulatory receptors | 0.69 | 0.48-0.89 | 0.040 |
| Other immune cells | B cells | 0.67 | 0.47-0.87 | 0.030 |
| T cells | Co-inhibitory receptprs | 0.67 | 0.48-0.86 | 0.031 |
| T cells | Immune effector cells | 0.66 | 0.47-0.86 | 0.031 |
| T cells | Th17 | 0.66 | 0.45-0.87 | 0.038 |
| T cells | Th1 | 0.65 | 0.45-0.85 | 0.030 |
| Other immune cells | Plasmacytoid dendritic cells | 0.64 | 0.43-0.85 | 0.030 |
| T cells | Naive T cells | 0.64 | 0.41-0.86 | 0.035 |
| T cells | Proliferating cytotoxic CD4 T cells | 0.63 | 0.42-0.84 | 0.030 |
| Other immune cells | Myeloid derived suppressor cells | 0.62 | 0.40-0.85 | 0.031 |
| Other immune cells | Myeloid dendritic cells | 0.62 | 0.41-0.84 | 0.030 |
| Other immune cells | NK cells | 0.62 | 0.42-0.82 | 0.030 |
| Stromal cells | Stromal signature | 0.62 | 0.41-0.83 | 0.030 |
| Stromal cells | Endothelial cells | 0.62 | 0.41-0.83 | 0.030 |
| T cells | Treg | 0.62 | 0.40-0.84 | 0.030 |
| Other immune cells | Macrophages | 0.60 | 0.39-0.81 | 0.030 |
| T cells | Adhesion molecules | 0.58 | 0.40-0.76 | 0.030 |
| Other immune cells | Neutrophil | 0.54 | 0.34-0.73 | 0.030 |

The receiver operating characteristic curves of individual signatures were compared against the TSE score applying the one-sided DeLong's test. P-values were adjusted using the Benjamini-Hochberg method.

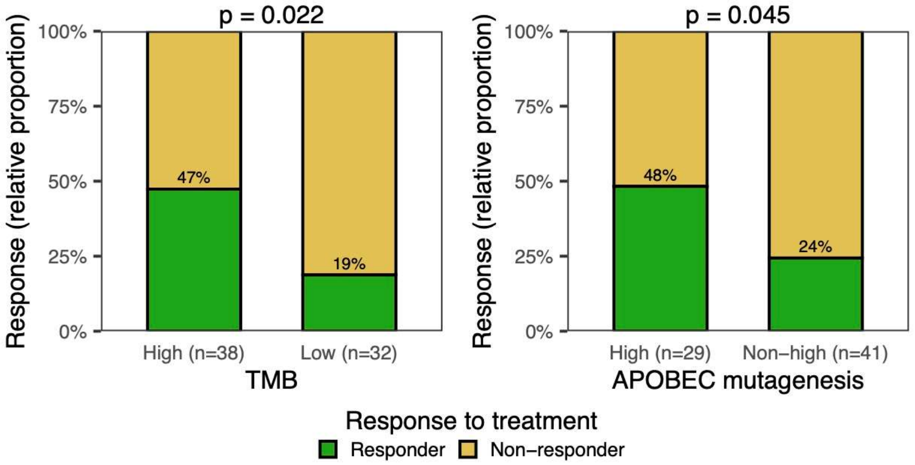


**Supplementary Figure 1. TMB and APOBEC mutagenesis in responders and non-responders to pembrolizumab**

Bar graphs display the number of responders and non-responders in patients with high and low tumor mutational burden (TMB), and in patients with high and non-high (medium, low or no) APOBEC mutagenesis. P-values were determined using the Fisher's exact test.

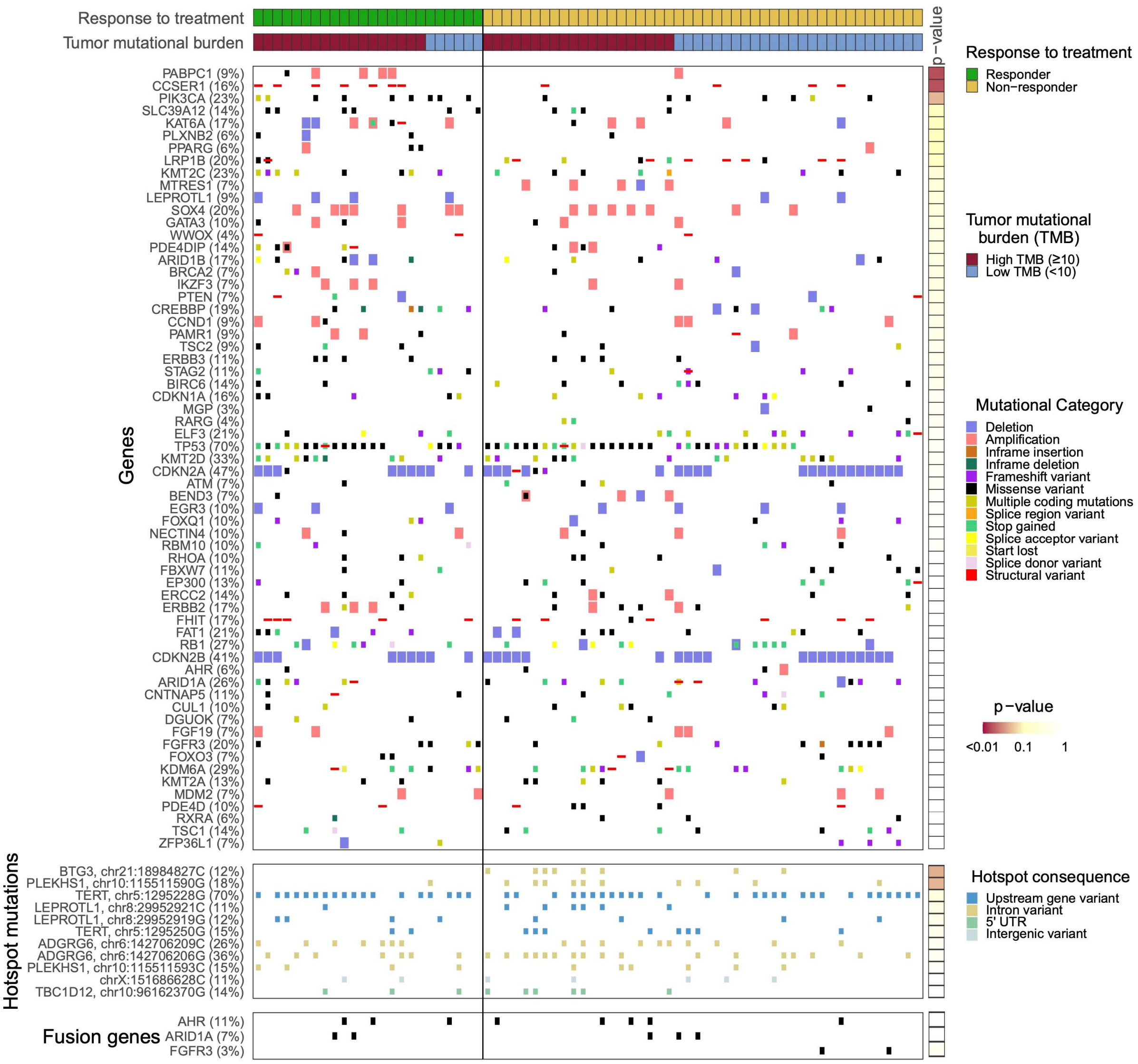


**Supplementary Figure 2. Gene alterations in responders and non-responders to pembrolizumab**

Overview of recurrent driver gene alterations, hotspot mutations and gene fusions in responders (n=24) and non-responders (n=46). Significantly mutated genes were estimated by dNdScv[9]; all genes with q < 0.05 were considered driver genes. Recurrent focal copy number changes were estimated by GISTIC2[10]; genes in genomic regions with q < 0.05 were considered significantly amplified/deleted. Gene fusions were detected from DNA using the pipeline described by the Hartwig Medical Foundation (https://github.com/hartwigmedical/). P-values of Fisher's exact test are shown on the right-hand side to reflect differential mutational status between responders and non-responders. After Benjamini-Hochberg correction, all p-values were not significant.

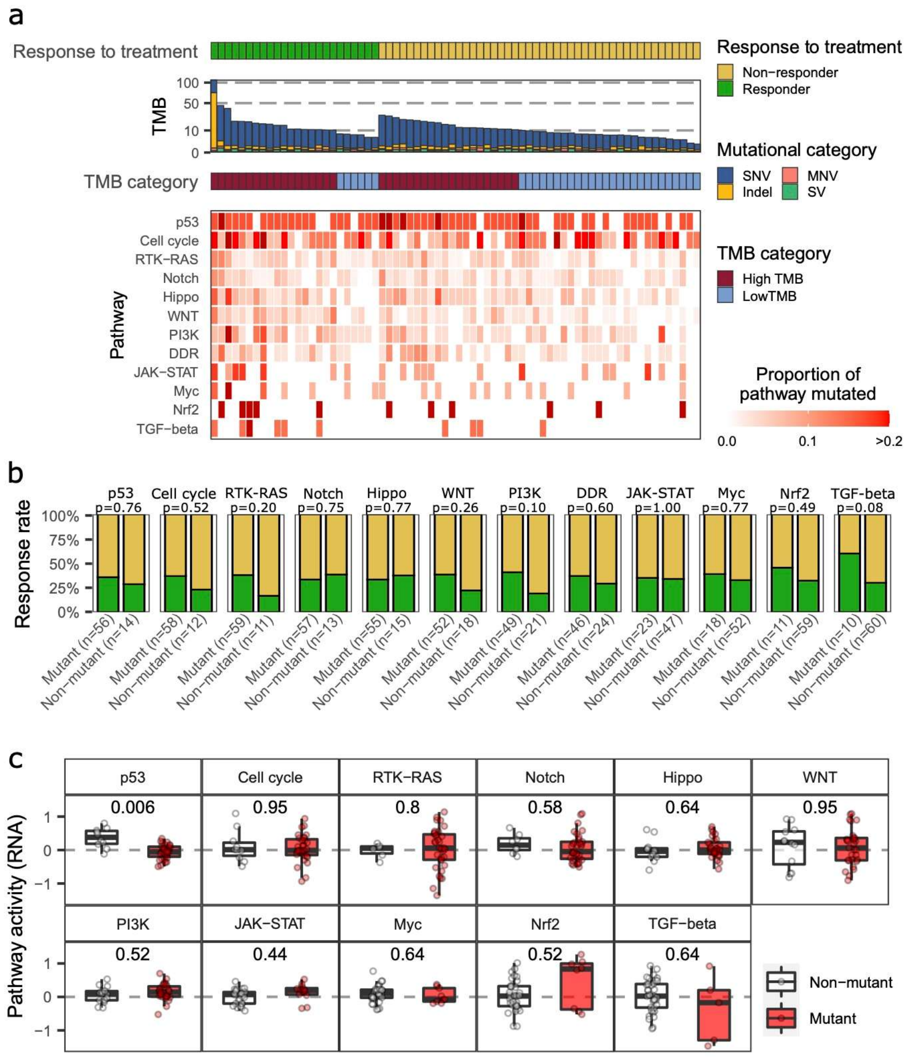


**Supplementary Figure 3. Pathway alterations in responders and non-responders to pembrolizumab**

**(a)** The proportion of altered genes per pathway is visualized by a gradient of red color. Genomic alterations include: non-synonymous mutations, structural variants and deep copy number changes (as defined by GISTIC). Genes included in this analysis were previously published[11-13] and are listed in Supplementary Data 1. Patients are stratified according to treatment response (responders, n=24; non-responders, n=46), and according to tumor mutational burden (TMB). (**b)** Proportion of responders and non-responders in samples with and without genomic pathway alterations. Differences between responders and non-responders were assessed for statistical significance using the Fisher's exact test. No correction for multiple testing was applied. (**c)** Pathway activity at RNA level in patients with genomic alterations in at least one gene (mutant) *versus* patients without any genomic alteration in a specific pathway (non-mutant). Wilcoxon-rank sum test with Benjamini-Hochberg correction was applied.

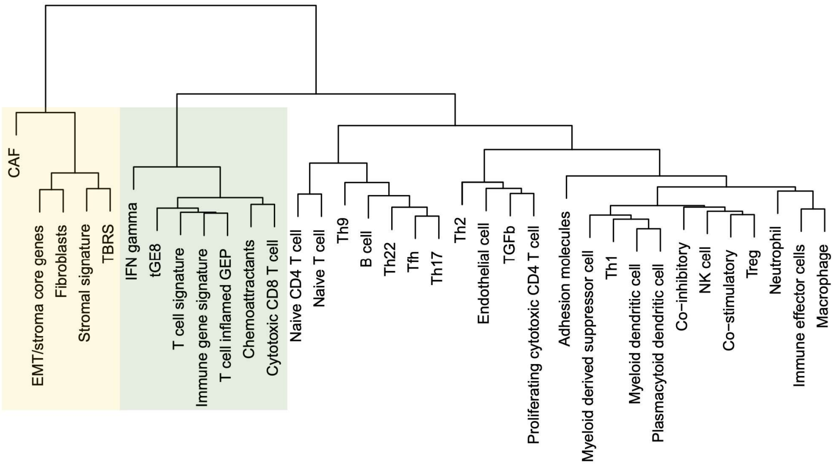


| Category | Signature | SD of score |
|---|---|---|
| Stromal resident cells | CAF | 1.31 |
| T cells | IFN gamma | 0.88 |
| Stromal resident cells | EMT/stroma core genes | 0.80 |
| Stromal resident cells | Fibroblasts | 0.79 |
| T cells | Naive CD4 T cell | 0.70 |
| T cells | Naive T cell | 0.69 |
| T cells | tGE8 | 0.67 |
| T cells | T cell signature | 0.62 |
| T cells | T cell inflamed GEP | 0.62 |
| T cells | Immune gene signature | 0.61 |
| T cells | Th9 | 0.60 |
| Stromal resident cells | TBRS | 0.54 |
| T cells | Cytotoxic CD8 T cell | 0.53 |
| Stromal resident cells | Stromal signature | 0.53 |
| Immune cells (non T cell) | Macrophage | 0.52 |
| Immune cells (non T cell) | B cell | 0.50 |
| T cells | Chemoattractants | 0.46 |
| T cells | Immune effector cells | 0.46 |
| Immune cells (non T cell) | Myeloid derived suppressor cell | 0.45 |
| T cells | Tfh | 0.43 |
| T cells | Th17 | 0.43 |
| Immune cells (non T cell) | Neutrophil | 0.42 |
| Immune cells (non T cell) | Plasmacytoid dendritic cell | 0.38 |
| T cells | Th22 | 0.37 |
| Immune cells (non T cell) | Myeloid dendritic cell | 0.37 |
| T cells | Co-inhibitory | 0.37 |
| T cells | Th1 | 0.36 |
| Immune cells (non T cell) | NK cell | 0.36 |
| T cells | Co-stimulatory | 0.34 |
| T cells | Adhesion molecules | 0.34 |
| Stromal resident cells | Endothelial cell | 0.34 |
| T cells | Treg | 0.33 |
| T cells | Th2 | 0.29 |
| Stromal resident cells | TGFb | 0.17 |
| T cells | Proliferating cytotoxic CD4 T cell | 0.12 |

**Supplementary Figure 4. Selection of gene signatures for T cells and stromal resident cells and their products towards usage in the TSE score**

The dendrogram represents the hierarchical clustering of gene signatures. The two groups with the highest standard deviations (SD) across all samples are highlighted. The table on the right shows all signatures and their standard deviation.

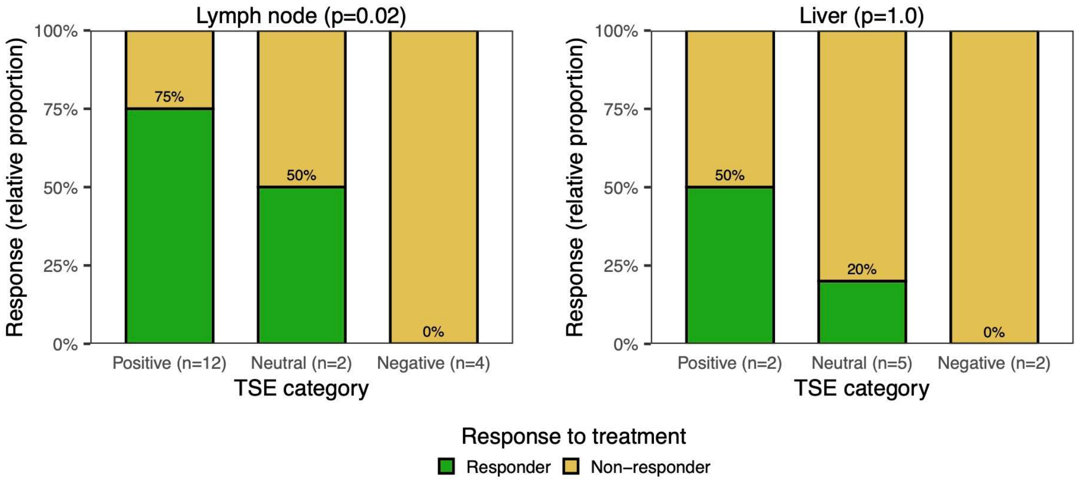


**Supplementary Figure 5. The TSE score predicts response to pembrolizumab regardless of metastatic site**

Bar graphs display the relative proportion of responders and non-responders in patients with a positive, neutral, or negative TSE score according to metastatic site. P-values of TSE positive *vs* negative were determined using the Fisher's exact test. A significant p-value was achieved in samples derived from lymph nodes, while the number of samples from liver was too limited to achieve any statistical significance.

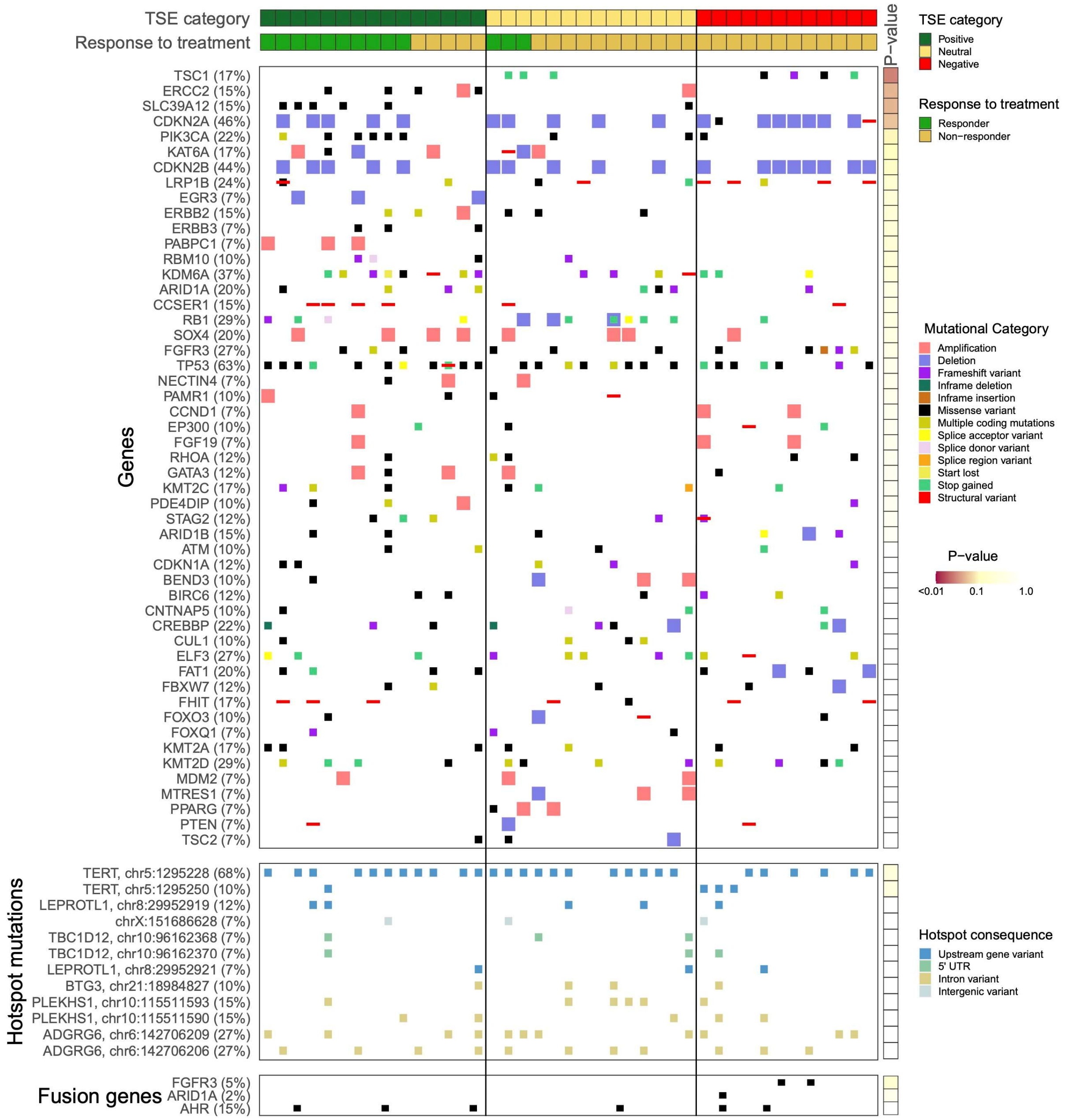


**Supplementary Figure 6. Gene alterations according to TSE score**

Recurrent driver gene alterations, hotspot mutations and gene fusions are displayed according to the T cell-to-stroma enrichment (TSE) score (TSE positive, n=15; TSE neutral, n=14; TSE negative, n=12). See legend to Supplementary Figure 2 for details. P-values of Fisher's exact test are shown on the right-hand side to reflect differential mutational status between patients with a positive and negative TSE score. No significant difference was observed after applying Benjamini-Hochberg correction for multiple testing.

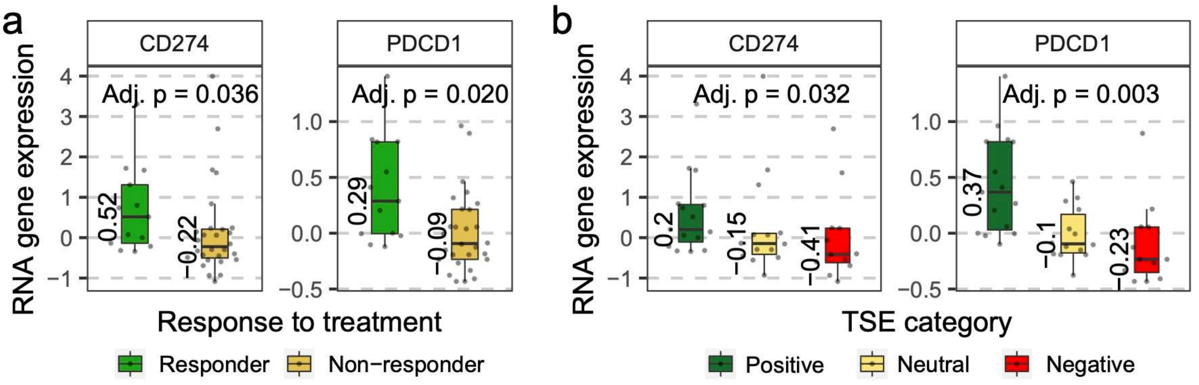


**Supplementary Figure 7. PD-1 and PD-L1 gene expression according to response to pembrolizumab and TSE score**

Boxplots displaying *CD274* (PD-L1) and *PDCD1* (PD-1) gene expression according to **(a)** response to pembrolizumab (responders, n=13; non-responders, n=28), or **(b)** the T cell-to-stroma enrichment (TSE) score (TSE positive, n=15; TSE neutral, n=14; TSE negative, n=12). Differences between response *vs* non-response or TSE positive *vs* negative were tested for statistical significance using the Wilcoxon-rank sum test and Benjamini-Hochberg corrected.

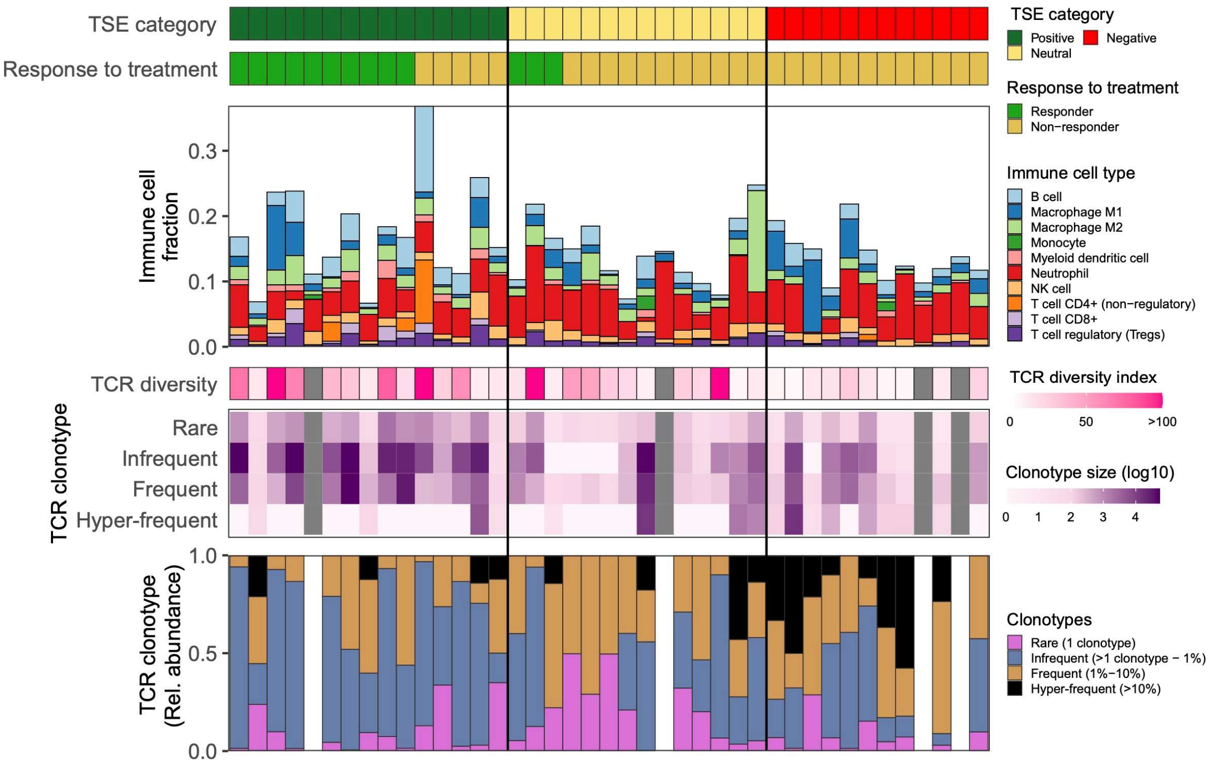


**Supplementary Figure 8. Immune cell fractions and T cell receptor repertoire characteristics according to TSE score**

Immune cell fractions and T cell receptor (TCR) repertoire characteristics are displayed according to T cell-to-stroma enrichment (TSE) score (TSE positive, n=15; TSE neutral, n=14; TSE negative, n=12). Immune cell fractions were determined using quanTIseq[14,15]. TCR diversity, as well as size and relative abundance of TCR clonotypes were determined using MiXCR v3.0.13[16].

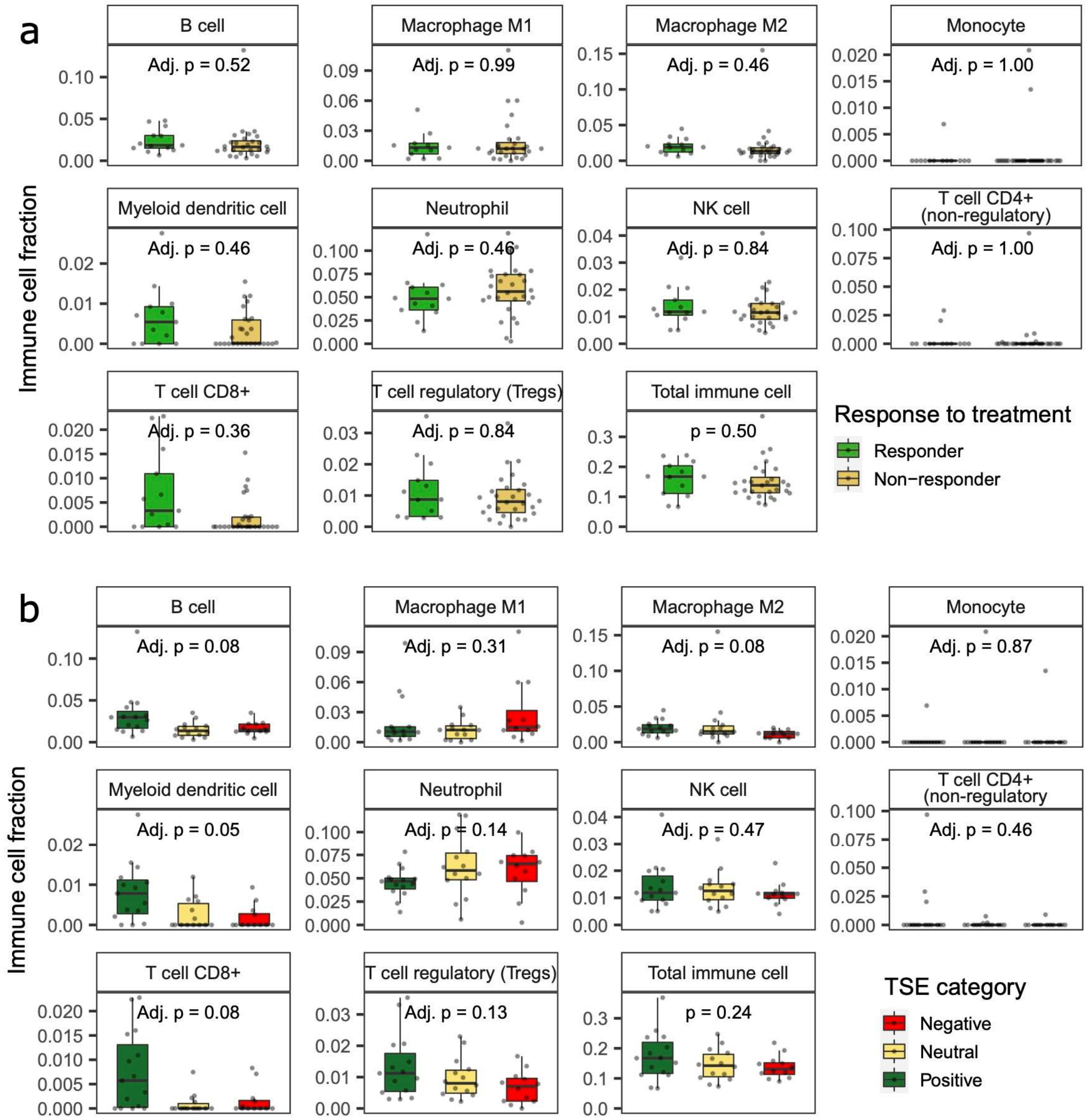


**Supplementary Figure 9. Relative abundance of individual immune cell populations according to response to pembrolizumab and TSE score**

**(a)** Boxplots displaying individual immune cell fractions according to response to pembrolizumab (responders, n=13; non-responders, n=28). (**b)** Boxplots displaying individual immune cell fractions according T cell-to-stroma enrichment (TSE) score (TSE positive, n=15; TSE neutral, n=14; TSE negative, n=12). Immune cell fractions were determined using quanTIseq[14,15]. Differences between response *vs* non-response or TSE positive *vs* negative were assessed for statistical significance using the Wilcoxon-rank sum test and Benjamini-Hochberg corrected.

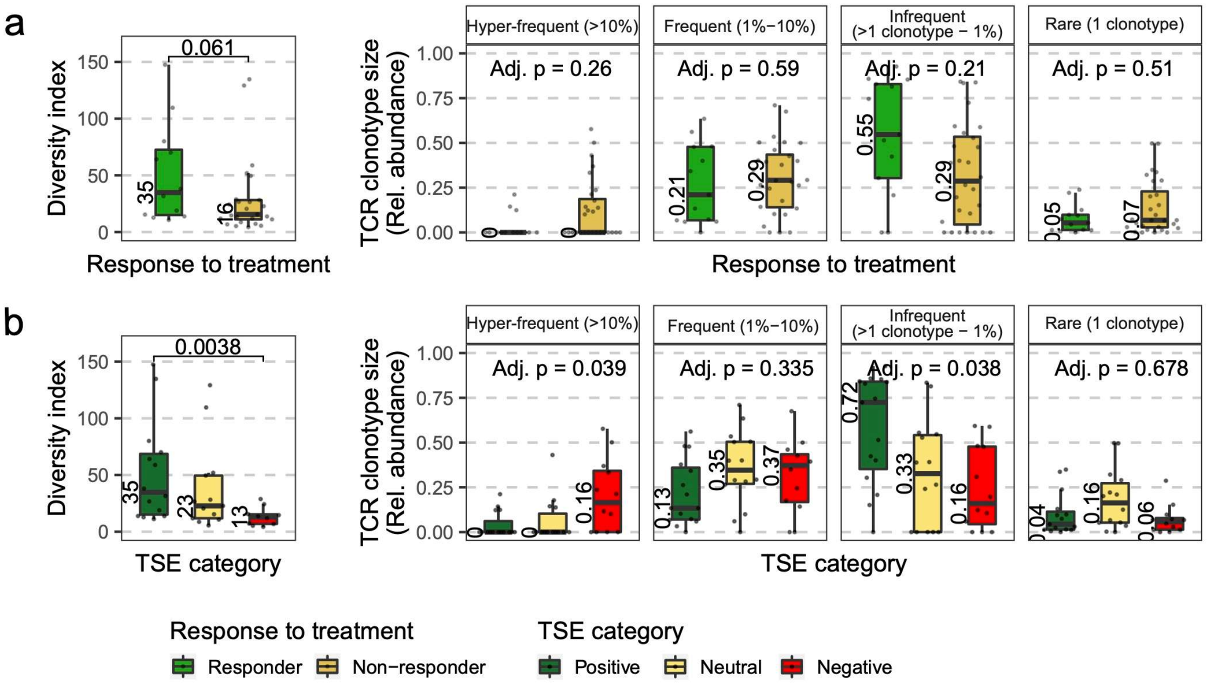


**Supplementary Figure 10. T cell receptor repertoire clonality and diversity according to response to pembrolizumab and TSE score**

Boxplots displaying TCR diversity and relative abundance of TCR clonotypes according to **(a)** response to pembrolizumab (responders, n=13; non-responders, n=28), or **(b)** T cell-to-stroma enrichment (TSE) score (TSE positive, n=15; TSE neutral, n=14; TSE negative, n=12). TCR diversity and clonality were determined using MiXCR v3.0.13[16]. Differences between response *vs* non-response or TSE positive *vs* negative were assessed for statistical significance using the Wilcoxon-rank sum test and Benjamini-Hochberg corrected.

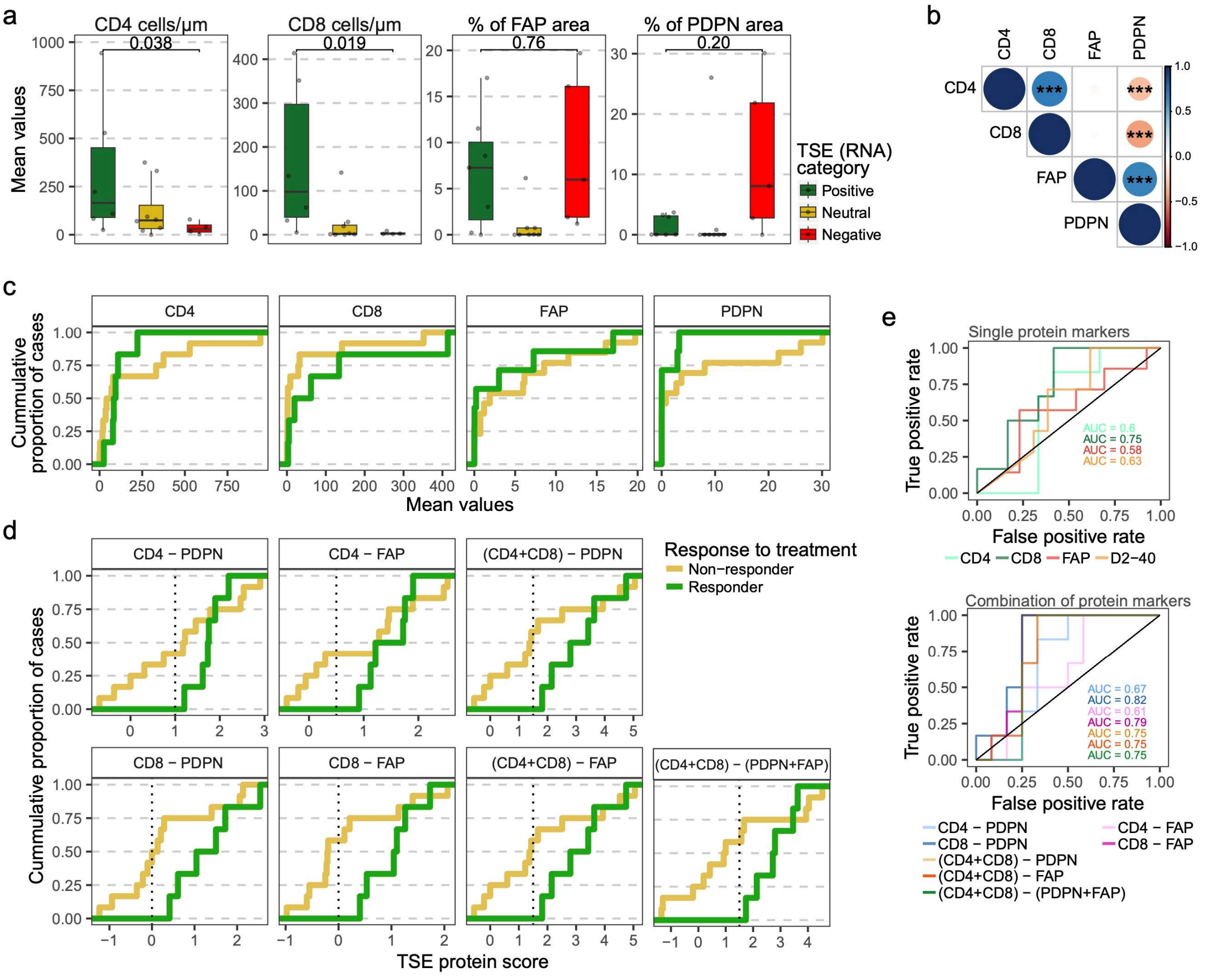


**Supplementary Figure 11. Individual and combinations of protein markers according to response to pembrolizumab and TSE score**

**(a)** Boxplots displaying densities of CD4 and CD8 T cells and percentages of marker-positive areas of of stromal resident cells and their products (FAP and PDPN) in tumors with a positive (n=7), neutral (n=8), or negative (n=5) T cell-to-stroma enrichment (TSE) score. Differential expression of these markers between TSE positive *vs* TSE negative was compared using the Wilcoxon-rank sum test (without correction for multiple testing). **(b)** Spearman correlation between protein markers (*** p<0.001). (**c**) Cumulative cases of responders and non-responders as a function of the individual protein marker values. (**d**) Performance of combinations of protein markers (TSE protein score) to select (~50% of) non-responders to pembrolizumab (indicated by a vertical dotted line). (**e**) Receiver operating characteristic (ROC) curves of individual protein markers and their combinations. The area under the curve (AUC) is displayed per condition.

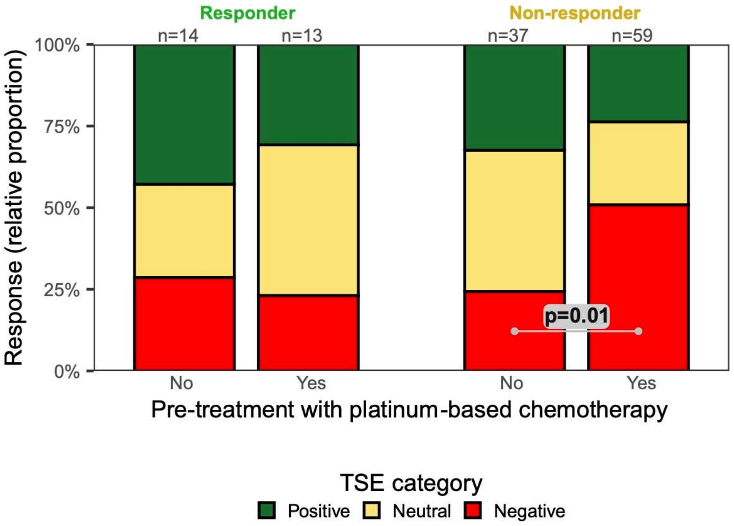


**Supplementary Figure 12. Effect of pretreatment on the TSE score according to response to atezolizumab in the IMvigor210 cohort**

Bar graphs display the relative proportion of tumors with a positive, neutral, or negative TSE score in responders and non-responders. Patients in the IMvigor210 cohort who only received platinum-based chemotherapy (excluding BCG pre-treated), were considered pre-treated. Patients were considered treatment-naïve when neither intravesical BCG nor platinum-based chemotherapy was previously administered. The Fisher exact test was applied to compare the proportion of TSE negative patients between pre-treated *vs* treatment-naive among non-responders.

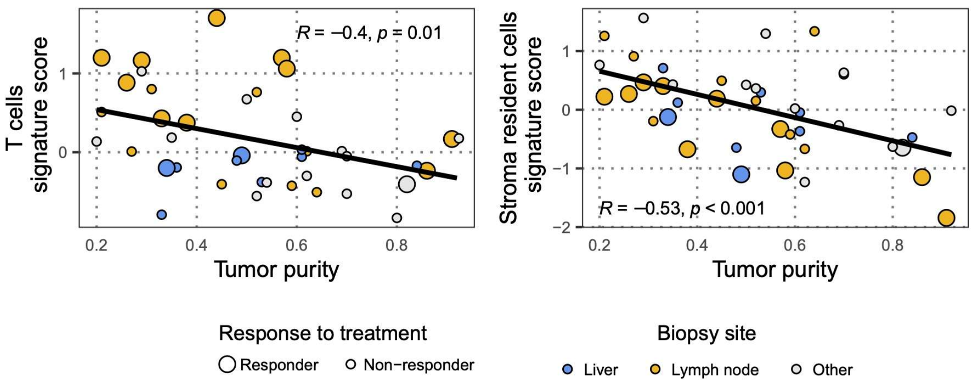


**Supplementary Figure 13. Signature scores representing T cells and stromal resident cells and products negatively correlate with tumor purity**

Plots display Pearson correlations between signature scores that are used to estimate the TSE score and tumor purity according to Pearson (left: T cells; right: stromal resident cells and products). Details regarding gene expression signatures and determination of tumor purity are described in Materials and Methods.

**Supplementary References**


1. Ayers M, Lunceford J, Nebozhyn M, et al: IFN-γ-related mRNA profile predicts clinical response to PD-1 blockade. J Clin Invest 127:2930-2940, 2017
2. Hammerl D, Massink MPG, Smid M, et al: Clonality, Antigen Recognition, and Suppression of CD8+ T Cells Differentially Affect Prognosis of Breast Cancer Subtypes. Clinical Cancer Research 26:505-517, 2020
3. Jerby-Arnon L, Shah P, Cuoco MS, et al: A Cancer Cell Program Promotes T Cell Exclusion and Resistance to Checkpoint Blockade. Cell 175:984-997.e24, 2018
4. Mariathasan S, Turley SJ, Nickles D, et al: TGFβ attenuates tumour response to PD-L1 blockade by contributing to exclusion of T cells. Nature 554:544-548, 2018
5. Oh DY, Kwek SS, Raju SS, et al: Intratumoral CD4(+) T Cells Mediate Anti-tumor Cytotoxicity in Human Bladder Cancer. Cell 181:1612-1625.e13, 2020
6. Spranger S, Bao R, Gajewski TF: Melanoma-intrinsic β-catenin signalling prevents anti-tumour immunity. Nature 523:231-235, 2015
7. Wang L, Saci A, Szabo PM, et al: EMT- and stroma-related gene expression and resistance to PD-1 blockade in urothelial cancer. Nature Communications 9:3503, 2018
8. Yoshihara K, Shahmoradgoli M, Martínez E, et al: Inferring tumour purity and stromal and immune cell admixture from expression data. Nature Communications 4:2612, 2013
9. Martincorena I, Raine KM, Gerstung M, et al: Universal Patterns of Selection in Cancer and Somatic Tissues. Cell 171:1029-1041.e21, 2017
10. Mermel CH, Schumacher SE, Hill B, et al: GISTIC2.0 facilitates sensitive and confident localization of the targets of focal somatic copy-number alteration in human cancers. Genome Biology 12:R41-R41, 2011
11. Sanchez-Vega F, Mina M, Armenia J, et al: Oncogenic Signaling Pathways in The Cancer Genome Atlas. Cell 173:321-337.e10, 2018
12. Leonard WJ: Role of JAK kinases and stats in cytokine signal transduction. International Journal of Hematology 73:271-277, 2001
13. Taber A, Christensen E, Lamy P, et al: Molecular correlates of cisplatin-based chemotherapy response in muscle invasive bladder cancer by integrated multi-omics analysis. Nature Communications 11:1-15, 2020
14. Sturm G, Finotello F, Petitprez F, et al: Comprehensive evaluation of transcriptome-based cell-type quantification methods for immuno-oncology. Bioinformatics 35:i436-i445, 2019
15. Finotello F, Mayer C, Plattner C, et al: Molecular and pharmacological modulators of the tumor immune contexture revealed by deconvolution of RNA-seq data. Genome Medicine 11:34-34, 2019
16. Bolotin DA, Poslavsky S, Mitrophanov I, et al: MiXCR: Software for comprehensive adaptive immunity profiling, Nature Publishing Group, 2015, pp 380-381
